# NCOR1 and OCT4 Facilitate Early Reprogramming by Co-Suppressing Fibroblast Gene Expression

**DOI:** 10.1101/747006

**Authors:** Georgina Peñalosa-Ruiz, Klaas W. Mulder, Gert Jan C. Veenstra

## Abstract

Reprogramming somatic cells to induced pluripotent stem cells (iPSC) succeeds only in a small fraction of cells within the population. Reprogramming occurs in distinctive stages, each facing its own bottlenecks. It initiates with overexpression of transcription factors OCT4, SOX2, KLF4 and c-MYC (OSKM) in somatic cells such as mouse embryonic fibroblasts (MEFs). OSKM bind chromatin, silencing the somatic identity and starting the stepwise reactivation of the pluripotency program. However, inefficient suppression of the somatic lineage leads to unwanted epigenetic memory from the tissue of origin, even in successfully generated iPSCs. Thus, it is essential to shed more light on chromatin regulators and processes involved in dissolving the somatic identity. Recent work characterized the role of transcriptional co-repressors NCOR1 and NCOR2 (also known as NCoR and SMRT), showing that they cooperate with c-MYC to silence pluripotency genes during late reprogramming stages. NCOR1/NCOR2 were also proposed to be involved in silencing fibroblast identity, however it is unclear how this happens. Here, we shed light on the role of NCOR1 in early reprogramming. We show that siRNA-mediated ablation of NCOR1 and OCT4 results in very similar phenotypes, including transcriptomic changes and highly correlated high content colony phenotypes.. Both NCOR1 and OCT4 bind to promoters co-occupied by c-MYC in MEFs. During early reprogramming, downregulation of one group of somatic MEF-expressed genes requires both NCOR1 and OCT4, whereas another group of MEF-expressed genes is downregulated by NCOR1 but not OCT4. Our data suggest that NCOR1, assisted by OCT4 and c-MYC, facilitates transcriptional inactivation of genes with high expression in MEFs, which need to be suppressed to bypass an early reprogramming block. This way, NCOR1 facilitates early reprogramming progression.

## INTRODUCTION

Induced pluripotent stem cells (iPSCs) are generated *in vitro* by overexpressing factors OCT4, SOX2, KLF4 and c-MYC (OSKM factors) in somatic cells [1]. iPSCs have been successfully used in disease modeling and cell transplantation, showcasing their relevance in regenerative medicine [2]. Despite significant improvements to the original protocol [3–5], in general, only a small percentage of somatic cells become pluripotent [6]. The reason is that, as embryonic development proceeds, cells gradually commit to a certain lineage, and acquire full stability upon differentiation [7]. Thus, *in vivo*, differentiated cells are not able to become other cell types, and such stability is conferred by chromatin state. Chromatin state defines the properties of regulatory elements as a function of DNA sequence, histone modifications, DNA methylation, chromatin architecture and the activity of transcription factor networks [7]. To understand reprogramming and develop new or improve existent iPS generation protocols, it is crucial to understand the interplay of chromatin modulators and transcriptional regulation.

Pluripotency acquisition involves silencing of somatic genes [8], and reactivation of the pluripotency network, which occurs in a stepwise manner [9]. The OSKM reprogramming transcription factors play different roles, but assist the two processes [10]. For instance, it has been shown that KLF4 and SOX2 bind to closed chromatin in MEFs [11,12]. These binding events are related to opening up of silenced pluripotency enhancers [12]. It has also been suggested that opening up of pluripotency enhancers requires cooperative binding of at least two of the reprogramming factors [10]. OCT4 was shown to bind to closed chromatin in somatic cells [11,13], but accumulating evidence indicates that OCT4 first binds open chromatin in MEFs and is recruited by other reprogramming factors to assist in opening-up of pluripotency enhancers [10,14,15]. OSK also are involved in transcriptional silencing of somatic genes, facilitating the relocation of somatic-specific transcription factors to loci co-occupied by OSK [10]. Thus, reprogramming factors facilitate the repression of the somatic program but also the activation of the pluripotency network. This transcriptional duality is elicited by differential interaction with transcriptional co-activators or co-repressors [16].

The transcriptional co-repressors NCOR1 (nuclear receptor co-repressor 1) and its paralogue NCOR2 (silencing mediator of retinoic acid and thyroid hormone receptor or NCoR2), interact with transcription factors to suppress their target genes [16]. NCOR1/NCOR2 are essential enzymatic co-factors of histone deacetylases, especially HDAC3 [17] and mediate transcriptional repression of their targets via deacetylation of Lysine residues of histone tails [18,19]. NCOR1/NCOR2 are essential for organism homeostasis, and have non-redundant roles in metazoans, as mutation of each gene individually leads to embryonic lethality [16]. In fact, NCOR1/NCOR2 regulate several essential metabolic and cellular processes, such as circadian rhythm [20] and mitochondrial function [21]. Recently, it was shown that NCOR1/NCOR2 interact with all four OSKM factors during reprogramming from MEFs to iPS [22]. NCOR1/NCOR2 interaction with c-MYC promotes the repression of pluripotency genes in later reprogramming stages [22], imposing a barrier for iPS generation. In addition, NCOR1/NCOR2 knockdowns induced transcriptional upregulation of somatic genes in the earliest reprogramming stages [22], arguing in favor of a dual role for these co-repressors. However, it is unknown how these early effects of NCOR1/SMART are brought about.

Previously, we performed a high-content imaging siRNA screening, combined with a secondary RNA-sequencing screen, to identify chromatin-associated regulators during early reprogramming from MEFs to iPS [23]. Using this data from two orthogonal screens, we identified strong phenotypic similarities between NCOR1 and OCT4 knockdowns. We have identified functional interactions from knockdowns showing similar phenotypes before [23,24]. Here we characterize the relationship of NCOR1 and OCT4 in early reprogramming. For such purpose, we first looked closer into the phenotypic similarities of both knockdowns, based on the high-content imaging. Then we set out to investigate the effect on the reprogramming transcriptome in *Ncor1* and *Oct4* knockdowns with RNA-sequencing. Finally, the analyses we compared our RNA-seq findings to published ChIP-seq and RNA-seq datasets. These analyses not only document the cooperation of OCT4 and NCOR1 in downregulating one set of somatic genes, they also show an antagonistic role in the regulation of another subset of somatic genes during early reprogramming.

## RESULTS

### High content screening reveals early reprogramming disruption upon *Ncor1* depletion

In a previous study, we performed a high-content siRNA screen, targeting 300 chromatin-associated factors during early reprogramming (**Fig. 1A**) [23]. We hypothesized that the onset of reprogramming would be associated with colony-level phenotypic changes and that these changes are affected upon gene perturbation. Moreover, using computational analysis of high-content imaging data, perturbed genes can be grouped according to their phenotypic similarities in a multidimensional space, rather than focusing on a single phenotype [25]. We used early pluripotency markers CDH1 and SALL4 to detect colonies at day 6 of reprogramming. Additionally, multiple colony features were extracted, including pluripotency marker expression levels, morphological traits such as colony roundness, area and symmetry, and many other colony texture and morphological features [23]. We selected 30 hits that were subjected to a secondary RNA-seq-based screening [23] (overview in **Fig. 1A**). We have previously identified functional interactions from knockdowns showing similar phenotypes [23,24,26]. Using this rationale, we compared knockdown-to-knockdown correlations based on high-content phenotypes and also based on transcriptomes (**Fig. 1B**). We filtered for pairwise correlations showing highest scores in both the high-content and the RNA-seq screen (both R> 0.4). We visualized the resulting potential interactions between genes (nodes) as networks, considering both high-content imaging correlations (edge width) and transcriptome-based correlations (edge color) of all siRNA knockdowns. We previously uncovered a functional interaction between BRCA1, BARD1 and WDR5 [23]. The strongest correlation however is between NCOR1-OCT4, raising the possibility of a functional interaction between these factors (**Fig. 1B**).

**Figure 1.**
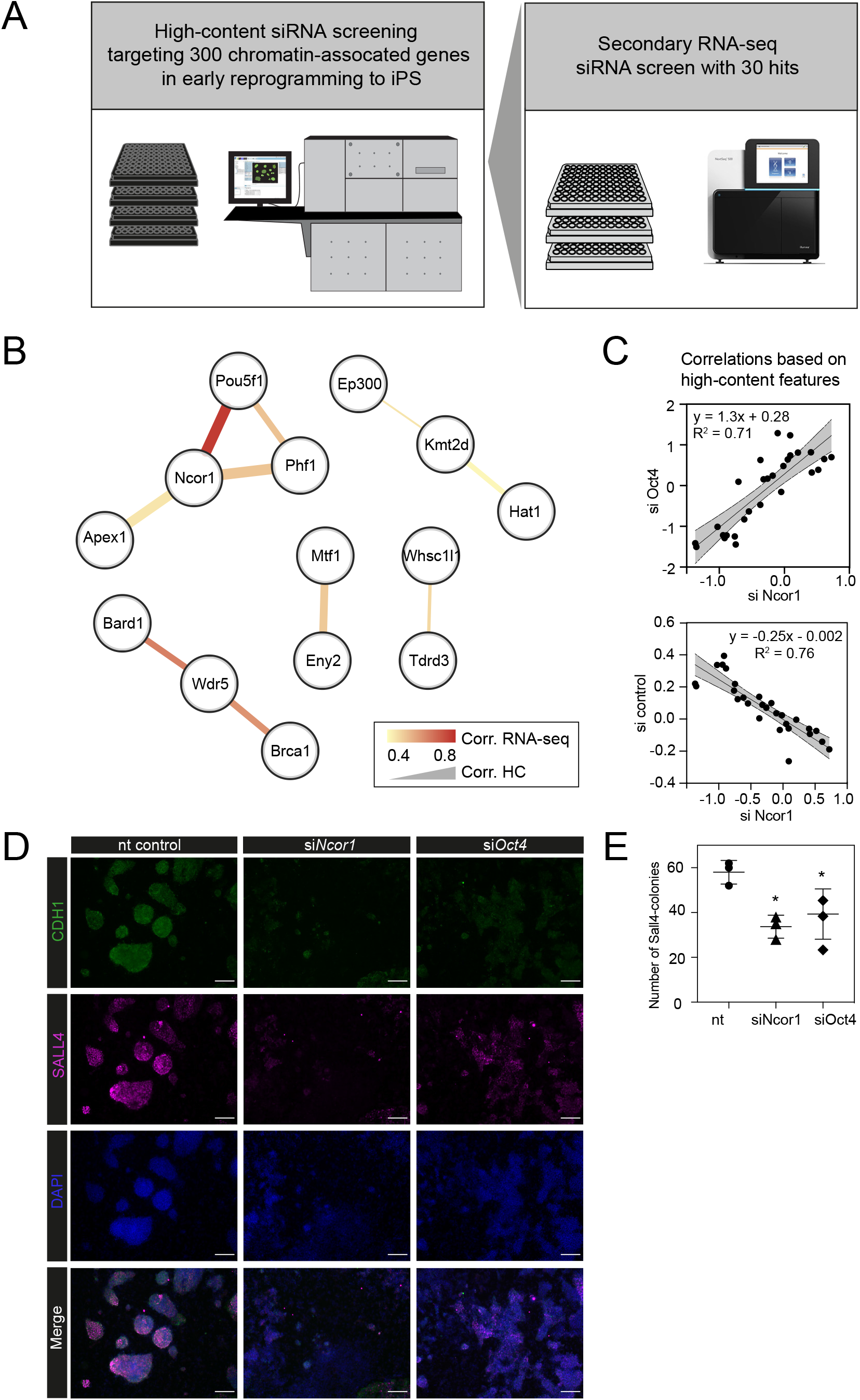
High content screening identifies Ncor1 as an early reprogramming modulator A) Overview of experimental setup for high-content siRNA screening in early reprogramming, followed by secondary RNA-seq-based screening, from which *Ncor1* was selected as candidate to follow-up. B) Gene network depicting correlations of high content colony phenotypes and RNA-seq profiles of genes targeted by siRNA. The nodes represent genes and the edges are pairwise Pearson correlations. The width of the edge represents correlation score by high-content imaging features, and the color of the edge represents the correlation strength based on RNA-sequencing. C) Scatterplots showing Pearson correlations according to 27 selected high-content features between *Ncor1* and *Oct4* knockdowns (upper panel) and between *Ncor1* and non-targeting (nt) control (lower panel). The middle line represents the best fit to linear regression model and the gray shades the 95 % confidence intervals (CI). The equation is the linear regression model and r2 is the correlation coefficient. D) Representative high-content images showing that, at day 6, *Ncor1* and *Oct4* depleted colonies display similar phenotypes, as compared to the non-targeting control. Scale bar represents 150 μM. In magenta nuclear transcription factor SALL4 and in green E-Cadherin (CDH1), counterstained with DAPI. E) Reprogramming efficiencies (Sall4-positive number of colonies) measured by incell western, compare *Ncor1* and *Oct4* knockdowns vs. nt control. *Ncor1* depletion impairs colony formation during early stages of reprogramming. Each data point represents one independent transfection and lines represent mean ± SD. Statistical significance analyzed by One-way ANOVA, (*) = p≤ 0.03.

We verified that, based on high-content features, *siNcor1* positively correlated with siOct4, but anti-correlated with the negative (nt, non-targeting) control (**Fig 1C**). Colony number analysis derived from the screen showed a significant reduction of colony formation upon *Ncor1* depletion (**Fig. 1D, Supplemental Fig.1**). High-content images from *Ncor1* and *Oct4* knockdowns showed very low CDH1 and SALL4 intensities and compromised colony formation, when compared to the nt control. We validated this finding using an independent In-Cell Western assay and confirmed that *siNcor1* and *siOct4* show similarly impaired reprogramming phenotypes (**Fig 1E, Supplemental Fig. 1**).

From these experiments we concluded that siRNA targeting of *Ncor1* results in an early reprogramming phenotype that is similar to that caused by *Oct4* knockdown.

### *Ncor1* depletion affects transcriptional regulation of fibroblast identity

To gain insight in the molecular events that explain early reprogramming disruption by *Ncor1* depletion, we performed gene expression analyses. We first asked whether *Ncor1* mRNA expression followed a dynamic pattern during early reprogramming. For this, *Ncor1* mRNA expression was quantified by RT-qPCR at different reprogramming timepoints, from MEFs to reprogramming day 6. We found that *Ncor1* mRNA expression was relatively stable throughout reprogramming, with slight variations (**Fig. 2A**). Then, we sought to understand the role of NCOR1 in transcriptional dynamics during early phases of iPSC reprogramming. For this purpose, we performed RNA-sequencing at day 3 and day 6 after expression of OSKM factors in MEFs transfected with control or *Ncor1* targeting siRNAs, respectively. Differential gene expression analysis revealed 405 deregulated genes at day 6 (siControl versus *siNcor1*, adjusted p-value <0.05, **Fig. 2B-C**). The effect size was relatively small for most transcripts, suggesting that the strong defects in repogramming observed with *siNcor1*, are brought about by moderate changes in expression of hundreds of transcripts (mean absolute log_2_-fold change of 0.43). The majority of the genes (94%) were up-regulated, as expected for a transcriptional co-repressor, although the deregulation of some of these genes may be an indirect consequence of the reduction of NCOR1 rather than a reduced NCOR1-mediated repression on that particular gene.

**Figure 2.**
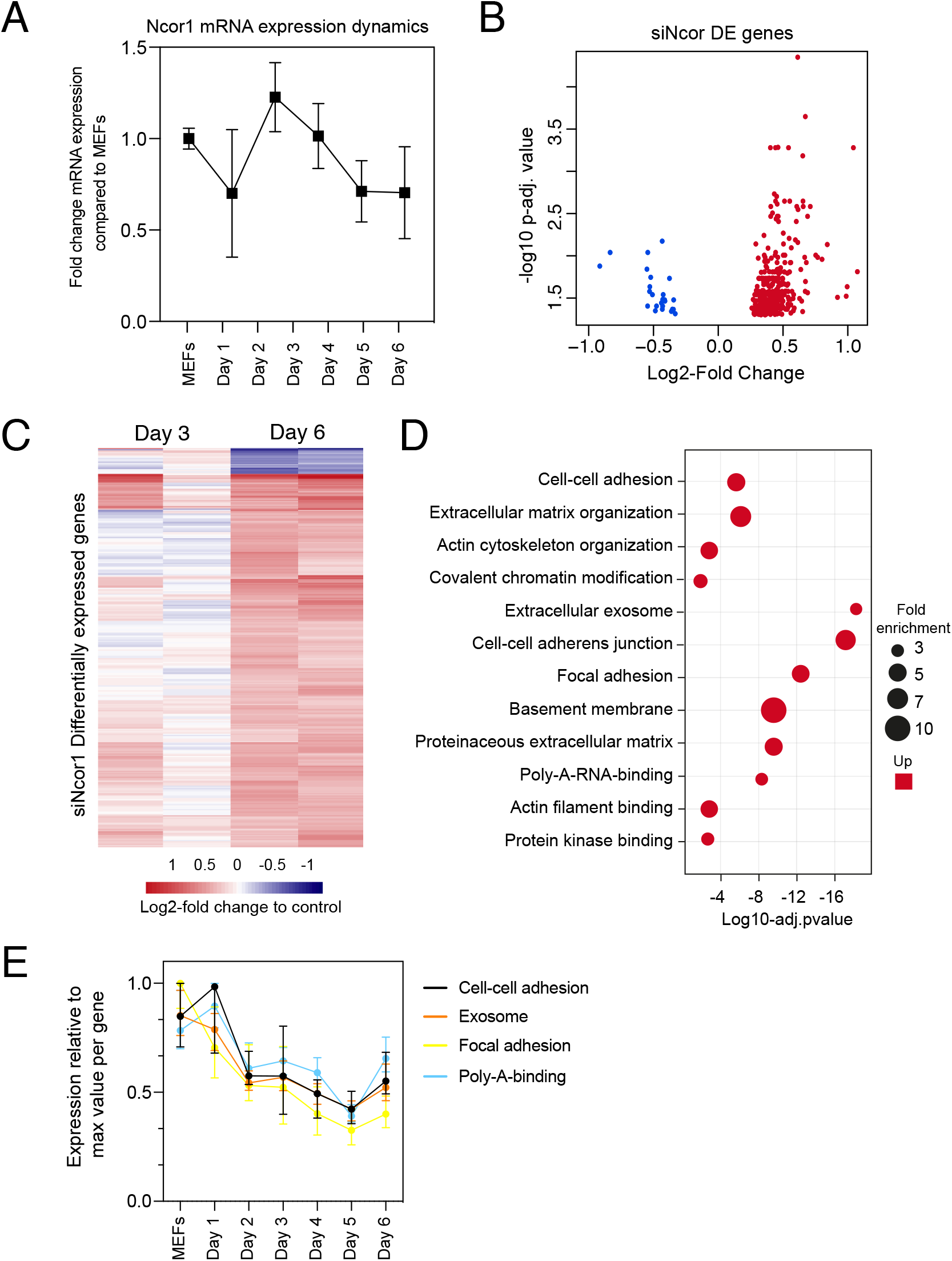
Gene-expression analysis of *Ncor1*-depleted reprogramming populations A) *Ncor1* mRNA expression dynamics during reprogramming time measured by RT-qPCR. Data are represented as fold change compared to MEFs (day 0). Data points represent the mean ± SD from two replicates from independent samples. B) Volcano plot for differentially expressed genes in *Ncor1* knockdown at reprogramming day 6, after RNA-sequencing. Blue data points represent the down-regulated genes and red datapoints represent up-regulated genes. C) Heatmap of differentially expressed genes from *siNcor1* at reprogramming day 3 and day 6. Color scale represents the log2-fold change compared to nontargeting control. D) Gene Ontology (GO) classification from differential genes in C represented as a bubble plot. The x-axis of the plot is the −log10 of the adjusted p-value, and some of the most significant terms are represented in red (upregulated). Size of the bubbles represents the fold-enrichment per category. E) Graphical representation of reprogramming RNA-seq temporal dynamics from some of the upregulated GO categories in D, shown in different colors. Each datapoint represents the median and the lines the 95% CI. Per GO category, medians were calculated based on the normalized expression value (log2-counts per million reads) of each gene, relative to the gene’s maximum value (maximum set to 1).

To gain insight into which processes were affected in *Ncor1*-depleted cells, we performed Gene Ontology classification (GO) for upregulated genes because the downregulated were very few (**Fig. 2D**). This analysis showed that upregulated genes were associated with cell-cell adhesion, poly-A-RNA-binding, exosome and cytoskeleton (**Fig. 2D**). Using our reprogramming RNA-seq time course dataset [23], we observed that genes associated with these terms were indeed downregulated in the course of early reprogramming (**Fig. 2E**). To note, the cell adhesion terms featured laminins, collagens and other genes related to basement membrane and extracellular matrix. These data suggest that reduction of *Ncor1* prevents downregulation of genes related to fibroblast identity during MEF to iPSC reprogramming.

### NCOR1 and OCT4 co-regulate two distinct MEF-expressed groups of genes

To gain insight into the functional relationship between OCT4 and NCOR1, we examined RNA-seq profiles at reprogramming day 6 for both knockdowns. To verify *Ncor1* and *Oct4* mRNA-targeting by the corresponding siRNAs, we plotted the RNA-seq normalized values for each of the knockdowns, compared to the controls at day 3 (**Supplemental Fig. 2**). *Oct4* knockdown did not show a high efficiency of knockdown at day 3 in these samples, however from time course experiments we have observed that *Oct4* mRNA is targeted and efficiently silenced at day 2, but rapidly recovers within a couple of days (**Supplemental Fig. 2**). This is due to its high overexpression driven by the tet-OSKM cassette [27]. Even with such transient early knockdown we observe a strong phenotype (Fig. 1) and many deregulated genes (Fig. 2A), highlighting the important contribution of OCT4 in OSKM reprogramming.

When we compared the genes deregulated in each knockdown we found that approximately half of the genes deregulated in *siNcor1* cells were also deregulated in *siOct4* cells (3.4 fold enrichment compared to random overlap, hypergeometric p-value 8.9 x 10^-54^; **Fig.3A**). After hierarchical clustering of genes deregulated by both *siNcor1* and *siOct4* based on RNA-seq, we identified one small cluster (Cluster 1) and a large cluster that was further divided in two (Cluster 2A, 2B; **Fig.3B**). Cluster 1 genes were relatively unaffected at day 3, but at day 6 they were downregulated for both knockdowns. For cluster 2A we found that genes were also unaffected at day 3, but showed a similar upregulated pattern in both *Oct4* and *Ncor1* siRNAs at day 6 (**Fig. 3B**). Cluster 2B contained genes with high expression in MEFs, most of which failed to be downregulated at day 3, whereas by day 6 they predominantly showed opposite expression changes in the knockdowns (*siNcor1* versus *siOct4*) (**Fig. 3B**). Our RNA-seq time course dataset [23] showed that genes in cluster 1 mostly remain the same in time, whereas clusters 2A and 2B tend to go down in time, but with different dynamics (**Fig. 3C**).

**Figure 3.**
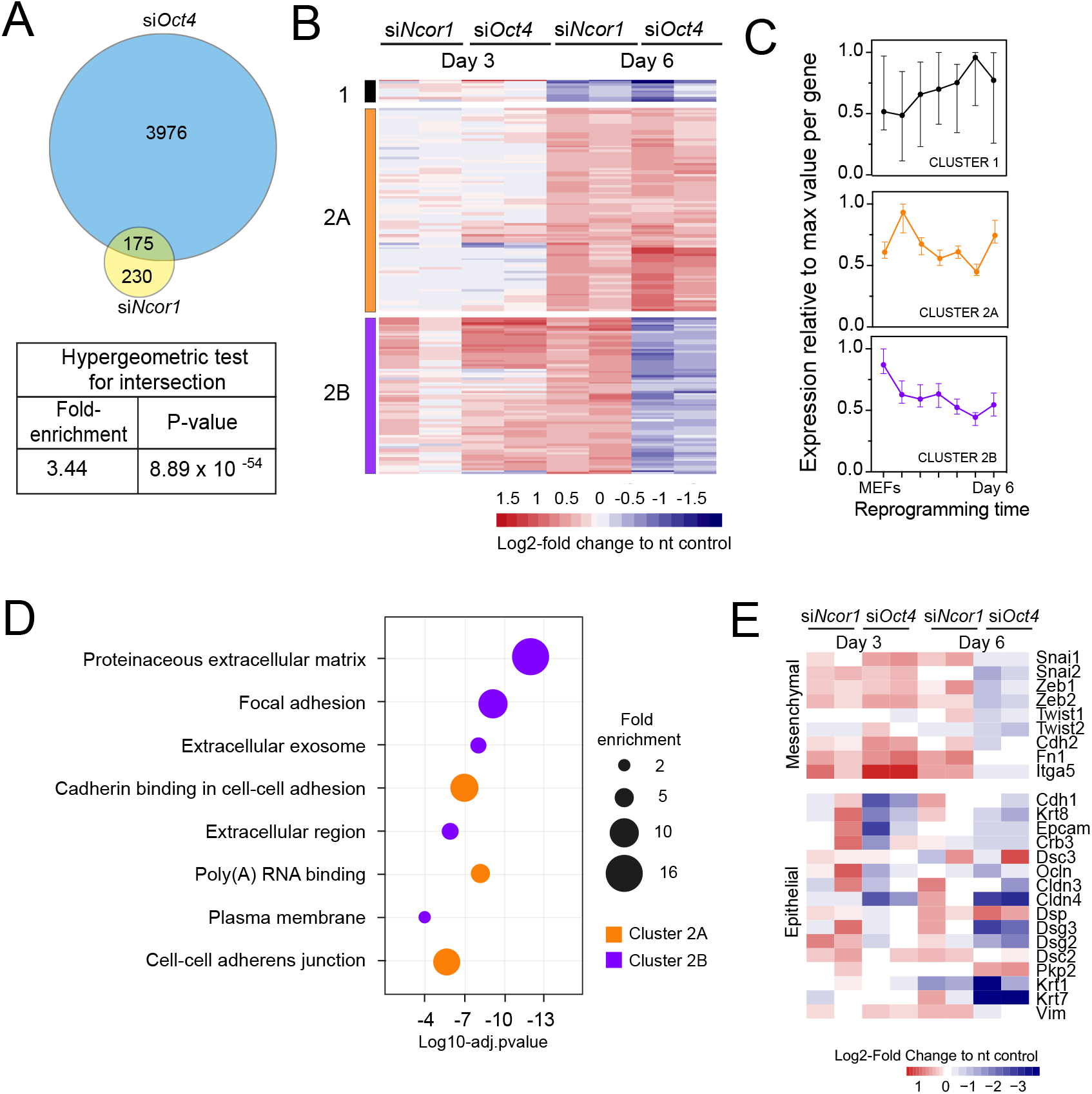
RNA-seq analysis shows that *Ncor1* and *Oct4* knockdowns show similar gene expression patterns. A) Venn-Euler diagram showing that *Ncor1* knockdown shares about half of deregulated genes with *Oct4* knockdown. The table underneath shows values for the hypergeometric test to show that the overlap of *Ncor1* and *Oct4* deregulated genes is significant. B) Heatmap of shared differential genes between *siNcor1* and *siOct4* from intersection in A, at reprogramming day 3 and day 6 for both siRNAs. Color scale represents the log2-fold change relative to non-targeting (nt) control. Clusters are tagged in three different colors, cluster 1 in black, cluster 2A in orange and cluster 2B in magenta C) Reprogramming RNA-seq temporal dynamics from clusters 1-3 shown in different colors, same color code as B. Each datapoint represents the median and the lines the 95 *%* CI. Per cluster, medians were calculated based on the normalized expression value (log2-counts per million reads) of each gene, relative to the gene’s maximum value = 1. D) Genes shown in clusters 1 and 2 from panel C were subjected to Gene Ontology (GO) classification. The bubble plot indicates some of the most significant classes. The size of the bubble represents the fold-enrichment and the −log10 of the adjusted p-value is shown on the x-axis. E) Heatmap representing expression values as log2-fold change ratio to control of mesenchymal and epithelial markers in *Oct4* and *Ncor1* knockdowns at reprogramming day 3 and day 6. The paired columns represent independent RNA-seq samples.

To shed some light on the difference between clusters 2A and 2B, we performed GO classification for genes of each cluster (**Fig. 3D**). NCOR1-OCT4 co-regulated genes (cluster 2A) showed an association with RNA metabolism and cell adhesion (**Fig. 3D**). The term “poly(A)-RNA binding” contained several components for pre-mRNA splicing. Genes increased by *siNcor1* but not *siOct4* (Cluster 2B), were associated with functions in extracellular matrix organization and other cell adhesion features. We wondered whether the “cell adhesion” terms were related to the mesenchymal-to-epithelial transition (MET) observed in early reprogramming [28,29]. We found that most mesenchymal genes exhibit a cluster 2B-type pattern, with increased expression at day 3 in both knockdowns and opposite effects of the two knockdowns at day 6. Epithelial genes on the other hand show more variable patterns (**Fig. 3E**). In *siOct4* cells, the downregulation of mesenchymal gene expression is delayed. By day 6, it not only catches up, this decrease in expression is more severe in *siOct4* cells relatively to control cells, suggesting that OCT4 moderates this decrease during normal reprogramming. In *siNcor1* cells on the other hand, mesenchymal gene expression is maintained at higher levels at both time points, although some epithelial genes are also expressed at higher levels (**Fig. 3E**).

These results identify two distinct groups of somatic genes, the expression of which is normally reduced during reprogramming. *Ncor1* depletion prevents downregulation of both groups, whereas *Oct4* depletion prevents the downregulation of one group and causes a stronger decrease in expression in the other group of genes.

### NCOR1 and OCT4 target promoters associated to cellular and metabolic processes in MEFs and early reprogramming cells

To gain insight into the functional relationship between NCOR1 and OCT4, we analyzed published ChIP-seq and RNA-seq data[10,22]. First, we asked whether the up-regulated genes that we identified in *siNcor1* and *siOct4* cells, were either MEF genes or genes dynamically changing during reprogramming. Consistent with our earlier analysis (**Fig. 3C**), these independent data confirmed that upregulated genes in *siNcor1* cells were mostly MEF-expressed and very early reprogramming (48h) genes, but were mostly inactive in ES cells (**Fig. 4A**). In the heatmap several genes related to fibroblast identity are labeled (lamins, collagens, fibronectin, signaling genes; Fig. 4A). In contrast, genes upregulated in *siOct4* cells showed a variety of expression patterns during reprogramming (**Fig. 4B**).

**Figure 4.**
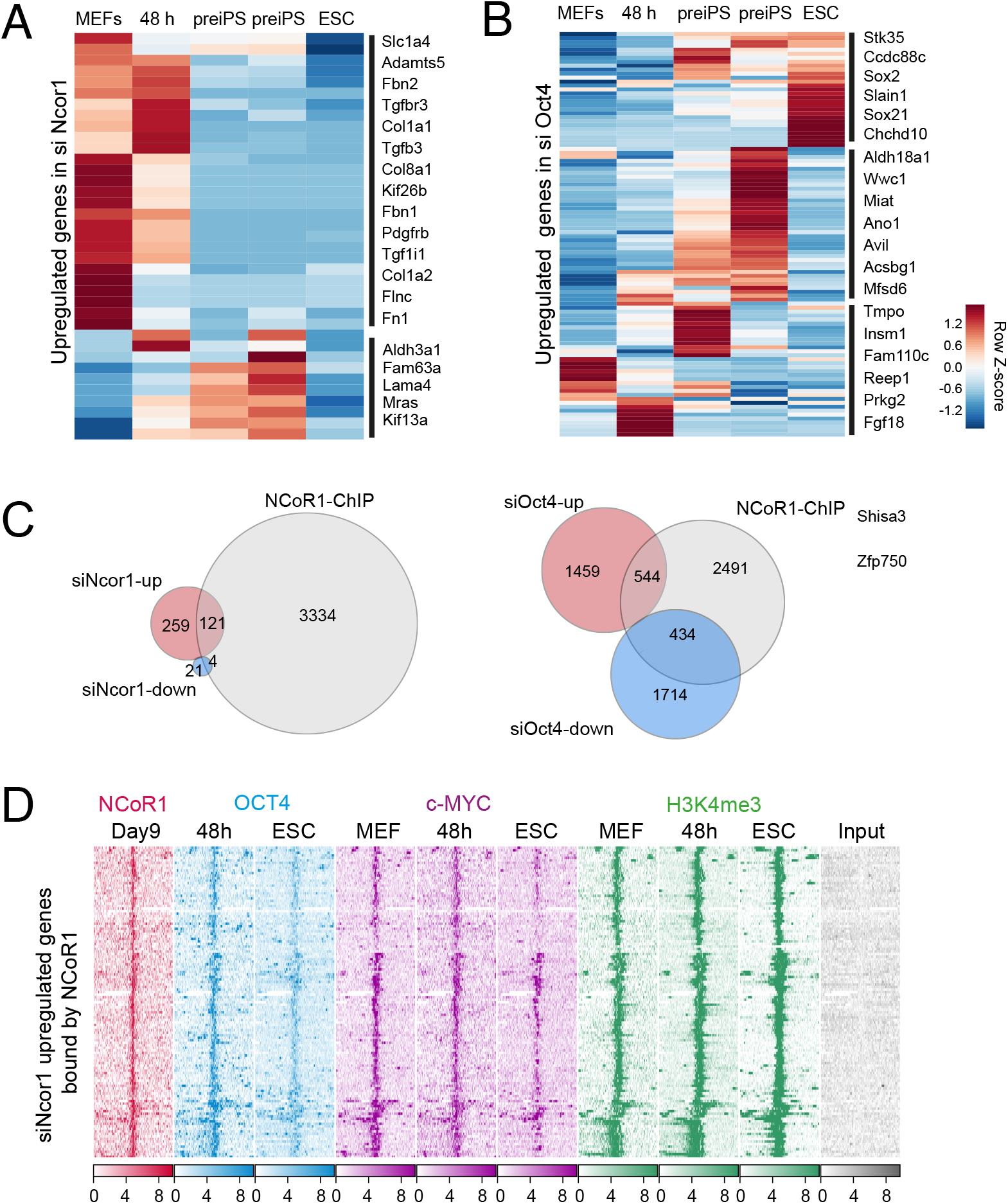
NCOR1 and OCT4 target promoters in MEFs and early reprogramming cells A) Expression of *siNcor1* up-regulated genes in MEFs, cells after 48 hours of reprogramming, pre-iPS intermediates and ESCs. Heatmap representation of differentially expressed genes during reprogramming [10] that are also upregulated by *siNcor1* in our study. The heatmap scale represents row-z scores of normalized expression values. B) Similar to panel A for *siOct4* upregulated genes in our study. Scales in both heatmaps are row Z-scores of normalized gene expression values. C) Venn-Euler diagram showing NCOR1 ChIP-seq peaks in reprogramming [22], overlapping with up-regulated or downregulated genes in *siNcor1* cells (left). Venn diagram showing NCOR1 ChIP-seq peaks in reprogramming overlapping with up-regulated or downregulated genes in *siOct4* (right). D) ChIP-seq enrichment of NCOR1, OCT4, c-MYC and the promoter-associated histone modification H3K4me3 [10,22] at NCOR/SMRT-bound genomic locations [22]) that are associated with *siNcor1*-upregulated genes in our data. The heatmap shows ChIP signals (color intensity) at genomic locations representing peak summits (center) plus and minus 5kb (left-right). Vertically different genomic locations are shown. Scales represent RPKM-normalized ChIP-seq signals.

Then, we asked which fraction of differentially expressed genes in *Ncor1-* or Oct4-knockdowns were directly bound by NCOR1. We overlapped NCOR1 binding sites in reprogramming cells with *siNcor1* and *siOct4* differential genes (**Fig.4C**, left and right, respectively) and calculated the fold enrichment. In all cases, the overlap was higher than expected by random chance, as determined with a hypergeometric test (**Table 1**). These data confirmed that NCOR1 directly binds to a significant fraction of deregulated genes in *Ncor1* and *Oct4* knockdowns (**Fig. 4C, Table 1**).

**Table 1.** Overlap of siNcor1 or siOct4 up/down regulated genes with NCOR1 ChIP-seq peaks. The overlapped was quantified as fold-enrichment and the statistical significance was calculated with a hypergeometric test.

| **Overlap comparison** | **Fold-Enrichment** | **p-value** |
| --- | --- | --- |
| siNcor1 up vs NCOR1 peaks | 3.2 | $5.9 \times 10^{-33}$ |
| siNcor1 down vs NCOR1 peaks | 1.6 | 0.096 |
| siOct4 up vs NCOR1 peaks | 2.7 | $5.3 \times 10^{-115}$ |
| siOct4 down vs NCOR1 peaks | 2.0 | $6.8 \times 10^{-50}$ |

We then asked whether NCOR1-bound genes with increased expression in *siNcor1* cells, were co-bound by OCT4 or c-MYC or were decorated with the H3K4me3 promoter mark (121 genes in **Fig. 4C left**). We observed that these NCOR1-bound regions were also enriched with OCT4 and c-MYC at 48 hours of reprogramming, whereas OCT4 and c-MYC binding was lower in ESC (**Fig.4D**). Since c-MYC is bound to those regions in MEFs already (**Fig. 4D**) and c-MYC, OCT4 and NCOR1 colocalize in the genome during reprogramming [22], it is possible that c-MYC facilitates OCT4 and NCOR1 binding to those regions early in reprogramming. Furthermore, the enrichment of H3K4me3 in MEFs, 48 h and ESC suggests that these genomic locations represent promoters (**Fig.4D**).

To explore the function of genes co-regulated by NCOR1 and OCT4, we performed a Gene Ontology classification of genes analyzed in Fig. 4D (**Supplemental Fig.3**). This analysis confirmed once more the terms we observed before (Figs. 2D and 3D), related to cell-cell adhesion, extracellular exosome, extracellular matrix, cytoskeleton, focal adhesion and poly(A) RNA binding, among others (Supplemental Fig.3). As before, the poly(A) RNA binding term included several genes related to translation and pre-mRNA splicing.

## DISCUSSION

In this study we found that depletion of *Ncor1* results in disrupted colony morphology in early iPS cells. Transcriptome analyses showed that NCOR1 has a role in suppression of fibroblast identity and signaling modulation (Fig. 2-3). Our findings extend an earlier report on the role of NCOR1 in later phases of reprogramming [22]. NCOR1 function could be mediated by c-MYC, that occupies many MEF regulatory regions [10].

Using multi-dimensional readouts, we found strong phenotypic similarities between *Ncor1* and *Oct4* depletion. Such phenotypic similarities often reflect functional interactions [23,24,26]. Our analyses indicate that NCOR1 and OCT4 co-bind a group of promoters that are also bound by c-MYC in early reprogramming cell populations. Genes associated with these promoters are expressed in MEFs and during the first 48 hours of reprogramming, and are mostly transcriptionally silent in ESCs (Fig.4). Upon *Ncor1* knockdown they are upregulated. This could mean that the repressor NCOR1 is recruited to such promoters by OCT4/c-MYC quite early in reprogramming, and transcriptional downregulation occurs via histone deacetylation. There was a surprisingly moderate effect on the transcriptome associated with *siNcor1*. Because of the increased proliferation of reprogramming cells [30], *Ncor1* knockdown at day 3 was 50%. Perhaps the effects on the transcriptome would have been stronger if we had transfected siRNAs at day 3 instead of day 0, or alternatively with a complete loss of function model. In addition, *Ncor1* and *Ncor2* may have partially redundant functions [31], which may have contributed to the relatively mild effect on the transcriptome. Nevertheless, we observe a strong phenotype in colony formation, and the transcriptome data suggest that *Ncor1* attenuates signaling, which might be crucial to weaken fibroblast identity. Consistent with this, NCOR1/NCOR2 are known to be connected to various signaling pathways [31].

Other genes involved are related to essential cellular functions, such as cell-cell adhesion, vesicle-mediated transport and RNA splicing. In agreement, complexes and proteins associated with these functions have been shown to be downregulated during intermediate stages of reprogramming [32]. Moreover some of these processes (e.g. cell adhesion, vesicle-mediated transport) have been shown to be barriers for reprogramming [33,34]. One apparent discrepancy in these analyses might be that the active promoter mark H3K4me3, seems abundant in MEFs, 48h reprogramming cells and ESCs. However, one possible explanation would be that target genes are downregulated in intermediate stages but then required again in late stages, as has been shown for some of the processes involved (cell adhesion, spliceosome) [32]. Moreover, some of these genes are essential for the cellular functions and continue to be expressed in ESCs, albeit at lower levels.

Recent work has shown that NCOR1/NCOR2 interact with all OSKM factors, but especially with c-MYC [22]. During late reprogramming stages, the co-repressors NCOR1/NCOR2 interact with c-MYC to silence the pluripotency network, posing a barrier for reprogramming [22]. Our study contributes with further characterization of an early facilitating function of NCOR1 in reprogramming, which in cooperation with OCT4 and c-MYC, may suppress cellular and metabolic functions sustaining fibroblast identity. This is consistent with a role of NCOR1 in overall cellular homeostasis and metabolic regulation [16,21]. It also highlights the emerging role and cooperation of reprogramming factors in transcriptional repression [10]. Future work will focus on potential functional interactions of NCOR1 with other chromatin factors, for instance with PHF1 or APEX1 (Fig.1B), and their physical interactions in early and late stages of reprogramming.

## DATA AVAILABILITY

High-content screening data can be downloaded from supplemental information in [23]. Reprogramming RNA-sequencing timecourse data is available from NCBI-GEO database accession number <u>GSE118679</u>. Non-targeting control RNA-seq data accession number in GEO is <u>GSE118677</u>.

## EXPERIMENTAL PROCEDURES

### Lentivirus production

HEK-293T cells up to passage 30 were cultured in 15 cm dishes to reach around 90 % confluency at the transfection day. Transfections with 3^rd^ generation lentiviruses were done with Lipofectamine 2000 following the manufacturer’s instructions. For each plate we used

### MEF-to-iPS Reprogramming

Primary mouse embryonic fibroblasts (MEFs) strain C57/BL6 passage 0-1 were seeded at a density of 10 000 cells per cm2 in MEF medium consisting of 15 % FBS, 100 nM b-mercaptoethanol and 1 % non-essential aminoacids (Thermo Scientific). Next day, MEFs were transduced with the doxycycline-inducible mouse tet-STEMCCA [27] and rtTA lentiviruses (Addgene # 20342) at an MOI of 1. One day after transduction, cells were either transfected with siRNAs or induced for reprogramming with reprogramming medium. This consisted of DMEM-high glucose (Thermo Scientific) supplemented with 10% stem cell grade FBS (Hyclone), 200 nM b-mercaptoethanol (Sigma), 1 % sodium pyruvate (Thermo Scientific), 2μg·mL-1 doxycycline, 3 μM Chiron (GSK3-inhibitor), 0.25 μM Alk5i (TGF-ß inhibitor), 50 μg·mL-1 ascorbic acid and 1·10^3^ U·mL-1 LIF. Next day after adding reprogramming medium was considered reprogramming Day 1.

### siRNA transfections

Transfections were performed as described previously [23]. Before adding the cell suspension containing OSKM-transduced MEFs, multi wells were prepared with transfection mix. This consisted of a 40nM pool of 3 different siRNAs per target mRNA, diluted in Optimem (Thermo Scientific) together with RNAiMAX lipofectamine, according to the manufacturers instructions. After this incubation, cell suspensions were added to each well. Twenty four hours after transfections, reprogramming started by adding medium with doxycycline.

### High-content screening

High-content screening was performed as described before[23]. Briefly, transfected cells were cultured in Cell Carrier 96-well black plates (Perkin Elmer), after six days, samples were fixed with 4 % PFA and stained for CDH1 (Cell Signaling 14472) and SALL4 (Abcam 29112) with DAPI counterstain. Plates were imaged in an Opera High-Content-Screening System (Perkin Elmer). Colony segmentation was done in multiple Z-planes using SALL4 staining. Columbus Software (Perkin Elmer) was used to extract all features in an automated fashion. Z-score normalization per plate was applied. The normalized data for high-content features were used for knockdown-to-knockdown Pearson-correlation analysis.

### RNA isolation and RT-qPCR

RNA was isolated with RNA Micro-prep kit (Zymo Research). Integrity was verified with Bioanalyzer and concentrations were determined with Nanodrop. For reverse transcription we used SuperScript III Kit (Thermo Scientific) starting with 120-180 ng total RNA. For the RT-qPCR reaction 1-2 ng of cDNA were used in 20 uL reaction mix with Sybr Green mix ready to use (iQ-SYBR-Green Supermix, Biorad). Relative gene expression was calculated with the ΔΔCt method using GAPDH as reference.

### RNA-sequencing library preparation and data analysis

Kapa-RNA HyperPrep kit with RiboErase (Roche, Kapa Biosystems) was used for ribosomal RNA depletion, and library preparation, with 200 ng total RNA. Library amplification was performed for 10 cycles, according to manufacturer’s instructions. Correct size distribution of 300 bp was verified with Bioanalyzer (Agilent Technologies). Quality controls were performed for selected transcripts in each sample before and after library preparation by qPCR.

NextSeq500 Illumina platform was used for paired-end library sequencing, with 43 bp read length. STAR v.2.5.b [35] was used to align reads to mouse genome assembly mm10. Data was normalized to log2-cpm with R package edgeR v.3.20.9 [36].

For differential gene expression analysis we used R Package DESeq2 v.1.18.1 [37]. Gene lists were filtered with a cutoff of p-adjusted value <0.05.

Gene Ontology classification with differential gene lists was performed with web-based DAVID [38], considering only categories with gene counts > 10 and p-adjusted value <0.05.

### ChIP-seq and RNA-seq data integration

For ChIP-seq data integration, FASTQ files of ChIP-seq data were downloaded from NCBI GEO (GSE70736, GSE90893, GSE90895) and mapped to the mm10 genome assembly using BWA allowing one mismatch per read. NCoR/SMRT peaks (Suppl. Table S4;[22]) were intersected with *siNcor1* day 6-upregulated genes (linux join, using gene symbols) to obtain genomic locations of interest. Heatmaps of ChIP-seq data were generated with fluff [39] with hierarchical clustering and RPKM normalization. RNA-seq data (Suppl. Table S2; [10]) was intersected with differentially expressed genes in our study (linux join, using gene symbols). Corresponding RNA-seq heatmaps were generated using seaborn clustermap with row clustering and row z-score normalization. Venn diagrams were obtained using venn3 from the matplotlib_venn library. Hypergeometric probabilities were calculated using scipy.stats.

## REFERENCES

[1] K. Takahashi, S. Yamanaka, Induction of pluripotent stem cells from mouse embryonic and adult fibroblast cultures by defined factors., Cell. 126 (2006) 663–76. doi:10.1016/j.cell.2006.07.024.

[2] H. Inoue, N. Nagata, H. Kurokawa, S. Yamanaka, iPS cells: a game changer for future medicine, EMBO J. 33 (2014) 409–417. doi: 10.1002/embj.201387098.

[3] M.A. Esteban, T. Wang, B. Qin, J. Yang, D. Qin, J. Cai, W. Li, Z. Weng, J. Chen, S. Ni, K. Chen, Y. Li, X. Liu, J. Xu, S. Zhang, F. Li, W. He, K. Labuda, Y. Song, A. Peterbauer, S. Wolbank, H. Redl, M. Zhong, D. Cai, L. Zeng, D. Pei, Vitamin C enhances the generation of mouse and human induced pluripotent stem cells., Cell Stem Cell. 6 (2010) 71–9. doi: 10.1016/j.stem.2009.12.001.

[4] S.E. Vidal, B. Amlani, T. Chen, A. Tsirigos, M. Stadtfeld, Combinatorial modulation of signaling pathways reveals cell-type-specific requirements for highly efficient and synchronous iPSC reprogramming, Stem Cell Reports. 3 (2014) 574–584. doi:10.1016/j.stemcr.2014.08.003.

[5] B. Di Stefano, S. Collombet, J.S. Jakobsen, M. Wierer, J.L. Sardina, A. Lackner, R. Stadhouders, C. Segura-Morales, M. Francesconi, F. Limone, M. Mann, B. Porse, D. Thieffry, T. Graf, C/EBPα creates elite cells for iPSC reprogramming by upregulating Klf4 and increasing the levels of Lsd1 and Brd4, Nat. Cell Biol. (2016). doi:10.1038/ncb3326.

[6] M. Stadtfeld, K. Hochedlinger, Induced pluripotency: History, mechanisms, and applications, Genes Dev. (2010). doi:10.1101/gad.1963910.

[7] M. Perino, G.J.C. Veenstra, Chromatin Control of Developmental Dynamics and Plasticity, Dev. Cell. 38 (2016) 610–620. doi:10.1016/J.DEVCEL.2016.08.004.

[8] J.M. Polo, E. Anderssen, R.M. Walsh, B.A. Schwarz, C.M. Nefzger, S.M. Lim, M. Borkent, E. Apostolou, S. Alaei, J. Cloutier, O. Bar-Nur, S. Cheloufi, M. Stadtfeld, M.E. Figueroa, D. Robinton, S. Natesan, A. Melnick, J. Zhu, S. Ramaswamy, K. Hochedlinger, A molecular roadmap of reprogramming somatic cells into iPS cells, Cell. 151 (2012) 1617–1632. doi: 10.1016/j.cell.2012.11.039.

[9] Y. Buganim, D.A. Faddah, A.W. Cheng, E. Itskovich, S. Markoulaki, K. Ganz, S.L. Klemm, A. Van Oudenaarden, R. Jaenisch, Single-cell expression analyses during cellular reprogramming reveal an early stochastic and a late hierarchic phase, Cell. 150 (2012) 1209–1222. doi:10.1016/j.cell.2012.08.023.

[10] C. Chronis, P. Fiziev, B. Papp, S. Butz, G. Bonora, S. Sabri, J. Ernst, K. Plath, Cooperative Binding of Transcription Factors Orchestrates Reprogramming, Cell. 168 (2017) 442–459.e20. doi: 10.1016/j.cell.2016.12.016.

[11] A. Soufi, M.F. Garcia, A. Jaroszewicz, N. Osman, M. Pellegrini, K.S. Zaret, Pioneer transcription factors target partial DNA motifs on nucleosomes to initiate reprogramming, Cell. 161 (2015) 555–568. doi:10.1016/j.cell.2015.03.017.

[12] D. Li, J. Liu, X. Yang, C. Zhou, J. Guo, C. Wu, Y. Qin, L. Guo, J. He, S. Yu, H. Liu, X. Wang, F. Wu, J. Kuang, A.P. Hutchins, J. Chen, D. Pei, Chromatin Accessibility Dynamics during iPSC Reprogramming, Cell Stem Cell. 21 (2017) 819–833.e6. doi:10.1016/j.stem.2017.10.012.

[13] A.S. Knaupp, S. Buckberry, J. Pflueger, S.M. Lim, E. Ford, M.R. Larcombe, F.J. Rossello, A. de Mendoza, S. Alaei, J. Firas, M.L. Holmes, S.S. Nair, S.J. Clark, C.M. Nefzger, R. Lister, J.M. Polo, Transient and Permanent Reconfiguration of Chromatin and Transcription Factor Occupancy Drive Reprogramming, Cell Stem Cell. 21 (2017) 834–845.e6. doi:10.1016/j.stem.2017.11.007.

[14] J. Donaghey, S. Thakurela, J. Charlton, J.S. Chen, Z.D. Smith, H. Gu, R. Pop, K. Clement, E.K. Stamenova, R. Karnik, D.R. Kelley, C.A. Gifford, D. Cacchiarelli, J.L. Rinn, A. Gnirke, M.J. Ziller, A. Meissner, Genetic determinants and epigenetic effects of pioneer-factor occupancy, Nat. Genet. 50 (2018) 250–258. doi:10.1038/s41588-017-0034-3.

[15] S. Xie, J. Duan, B. Li, P. Zhou, G.C. Hon, Multiplexed Engineering and Analysis of Combinatorial Enhancer Activity in Single Cells., Mol. Cell. 66 (2017) 285–299.e5. doi:10.1016/j.molcel.2017.03.007.

[16] A. Mottis, L. Mouchiroud, J. Auwerx, Emerging roles of the corepressors NCoR1 and SMRT in homeostasis, Genes Dev. 27 (2013) 819–835. doi:10.1101/gad.214023.113.

[17] S.H. You, H.W. Lim, Z. Sun, M. Broache, K.J. Won, M.A. Lazar, Nuclear receptor co-repressors are required for the histone-deacetylase activity of HDAC3 in vivo, Nat. Struct. Mol. Biol. 20 (2013) 182–187. doi: 10.1038/nsmb.2476.

[18] L. Alland, R. Muhle, H. Hou, J. Potes, L. Chin, N. Schreiber-Agus, R.A. DePinho, Role for N-CoR and histone deacetylase in Sin3-mediated transcriptional repression, Nature. 387 (1997) 49–55. doi: 10.1038/387049a0.

[19] T. Heinzel, R.M. Lavinsky, T.-M. Mullen, M. Söderström, C.D. Laherty, J. Torchia, W.-M. Yang, G. Brard, S.D. Ngo, J.R. Davie, E. Seto, R.N. Eisenman, D.W. Rose, C.K. Glass, M.G. Rosenfeld, A complex containing N-CoR, mSln3 and histone deacetylase mediates transcriptional repression, Nature. 387 (1997) 43–48. doi:10.1038/387043a0.

[20] T. Alenghat, K. Meyers, S.E. Mullican, K. Leitner, A. Adeniji-Adele, J. Avila, M. Bućan, R.S. Ahima, K.H. Kaestner, M.A. Lazar, Nuclear receptor corepressor and histone deacetylase 3 govern circadian metabolic physiology, Nature. 456 (2008) 997–1000. doi:10.1038/nature07541.

[21] W. Fan, R. Evans, PPARs and ERRs: molecular mediators of mitochondrial metabolism, Curr. Opin. Cell Biol. 33 (2015) 49–54. doi:10.1016/J.CEB.2014.11.002.

[22] Q. Zhuang, W. Li, C. Benda, Z. Huang, T. Ahmed, P. Liu, X. Guo, D.P. Ibañez, Z. Luo, M. Zhang, M.M. Abdul, Z. Yang, J. Yang, Y. Huang, H. Zhang, D. Huang, J. Zhou, X. Zhong, X. Zhu, X. Fu, W. Fan, Y. Liu, Y. Xu, C. Ward, M.J. Khan, S. Kanwal, B. Mirza, M.D. Tortorella, H.F. Tse, J. Chen, B. Qin, X. Bao, S. Gao, A.P. Hutchins, M.A. Esteban, NCoR/SMRT co-repressors cooperate with c-MYC to create an epigenetic barrier to somatic cell reprogramming, Nat. Cell Biol. (2018). doi:10.1038/s41556-018-0047-x.

[23] G. Peñalosa-Ruiz, V. Bousgouni, J.P. Gerlach, S. Waarlo, J. V. van de Ven, T.E. Veenstra, J.C.R. Silva, S.J. van Heeringen, C. Bakal, K.W. Mulder, G.J.C. Veenstra, WDR5, BRCA1, and BARD1 Co-regulate the DNA Damage Response and Modulate the Mesenchymal-to-Epithelial Transition during Early Reprogramming, Stem Cell Reports. 12 (2019) 743–756. doi: 10.1016/j.stemcr.2019.02.006.

[24] K.W. Mulder, X. Wang, C. Escriu, Y. Ito, R.F. Schwarz, J. Gillis, G. Sirokmány, G. Donati, S. Uribe-Lewis, P. Pavlidis, A. Murrell, F. Markowetz, F.M. Watt, Diverse epigenetic strategies interact to control epidermal differentiation., Nat. Cell Biol. 14 (2012) 753–63. doi: 10.1038/ncb2520.

[25] M. Boutros, L.P. Brás, W. Huber, Analysis of cell-based RNAi screens, Genome Biol. 7 (2006). doi:10.1186/gb-2006-7-7-r66.

[26] S.E.J. Tanis, P.W.T.C. Jansen, H. Zhou, S.J. van Heeringen, M. Vermeulen, M. Kretz, K.W. Mulder, Splicing and Chromatin Factors Jointly Regulate Epidermal Differentiation, Cell Rep. 25 (2018) 1292–1303.e5. doi:10.1016/J.CELREP.2018.10.017.

[27] C. a Sommer, M. Stadtfeld, G.J. Murphy, K. Hochedlinger, D.N. Kotton, G. Mostoslavsky, Induced pluripotent stem cell generation using a single lentiviral stem cell cassette., Stem Cells. 27 (2009) 543–9. doi: 10.1634/stemcells.2008-1075.

[28] R. Li, J. Liang, S. Ni, T. Zhou, X. Qing, H. Li, W. He, J. Chen, F. Li, Q. Zhuang, B. Qin, J. Xu, W. Li, J. Yang, Y. Gan, D. Qin, S. Feng, H. Song, D. Yang, B. Zhang, L. Zeng, L. Lai, M.A. Esteban, D. Pei, A mesenchymal-to-Epithelial transition initiates and is required for the nuclear reprogramming of mouse fibroblasts, Cell Stem Cell. 7 (2010) 51–63. doi:10.1016/j.stem.2010.04.014.

[29] P. Samavarchi-Tehrani, A. Golipour, L. David, H.K. Sung, T.A. Beyer, A. Datti, K. Woltjen, A. Nagy, J.L. Wrana, Functional genomics reveals a BMP-Driven mesenchymal-to-Epithelial transition in the initiation of somatic cell reprogramming, Cell Stem Cell. 7 (2010) 64–77. doi:10.1016/j.stem.2010.04.015.

[30] S. Ruiz, A.D. Panopoulos, A. Herrerías, K.-D. Bissig, M. Lutz, W.T. Berggren, I.M. Verma, J.C. Izpisua Belmonte, A High Proliferation Rate Is Required for Cell Reprogramming and Maintenance of Human Embryonic Stem Cell Identity, Curr. Biol. 21 (2011) 45–52. doi:10.1016/J.CUB.2010.11.049.

[31] V. Perissi, K. Jepsen, C.K. Glass, M.G. Rosenfeld, Deconstructing repression: evolving models of co-repressor action, Nat. Rev. Genet. 11 (2010). doi: 10.1038/nrg2736.

[32] J. Hansson, M.R. Rafiee, S. Reiland, J.M. Polo, J. Gehring, S. Okawa, W. Huber, K. Hochedlinger, J. Krijgsveld, Highly Coordinated Proteome Dynamics during Reprogramming of Somatic Cells to Pluripotency, Cell Rep. 2 (2012) 1579–1592. doi:10.1016/j.celrep.2012.10.014.

[33] S.M. Buckley, B. Aranda-Orgilles, A. Strikoudis, E. Apostolou, E. Loizou, K. Moran-Crusio, C.L. Farnsworth, A.A. Koller, R. Dasgupta, J.C. Silva, M. Stadtfeld, K. Hochedlinger, E.I. Chen, I. Aifantis, Regulation of Pluripotency and Cellular Reprogramming by the Ubiquitin-Proteasome System, Cell Stem Cell. 11 (2012) 783–798. doi:10.1016/J.STEM.2012.09.011.

[34] H. Qin, A. Diaz, L. Blouin, R.J. Lebbink, W. Patena, P. Tanbun, E.M. LeProust, M.T. McManus, J.S. Song, M. Ramalho-Santos, Systematic Identification of Barriers to Human iPSC Generation, Cell. 158 (2014) 449–461. doi: 10.1016/j.cell.2014.05.040.

[35] A. Dobin, C.A. Davis, F. Schlesinger, J. Drenkow, C. Zaleski, S. Jha, P. Batut, M. Chaisson, T.R. Gingeras, STAR: ultrafast universal RNA-seq aligner, Bioinformatics. 29 (2013) 15–21. doi: 10.1093/bioinformatics/bts635.

[36] M.D. Robinson, D.J. McCarthy, G.K. Smyth, edgeR: a Bioconductor package for differential expression analysis of digital gene expression data, Bioinformatics. 26 (2010) 139–140. doi:10.1093/bioinformatics/btp616.

[37] M.I. Love, W. Huber, S. Anders, Moderated estimation of fold change and dispersion for RNA-seq data with DESeq2, Genome Biol. 15 (2014) 550. doi: 10.1186/s13059-014-0550-8.

[38] D.W. Huang, B.T. Sherman, R.A. Lempicki, Systematic and integrative analysis of large gene lists using DAVID bioinformatics resources, Nat. Protoc. 4 (2009) 44–57. doi:10.1038/nprot.2008.211.

[39] G. Georgiou, S.J. van Heeringen, fluff: exploratory analysis and visualization of high-throughput sequencing data, PeerJ. 4 (2016) e2209. doi: 10.7717/peerj.2209.

