## Supplemental Information for "NCOR1 and OCT4 Facilitate Early Reprogramming by Co-Suppressing Fibroblast Gene Expression"

A

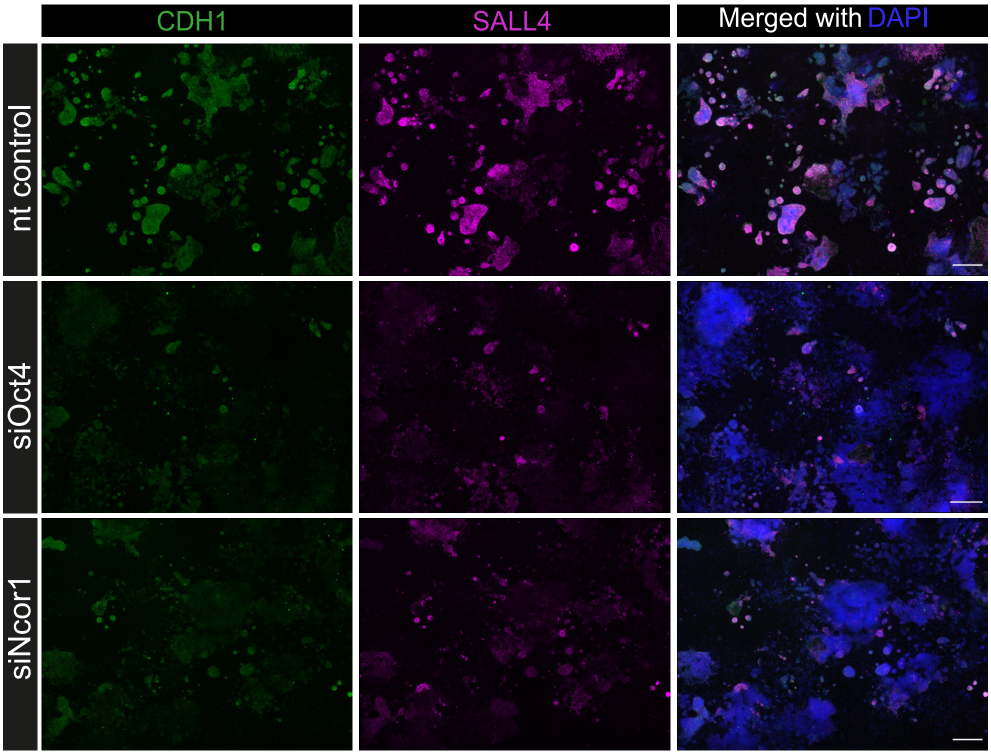


B

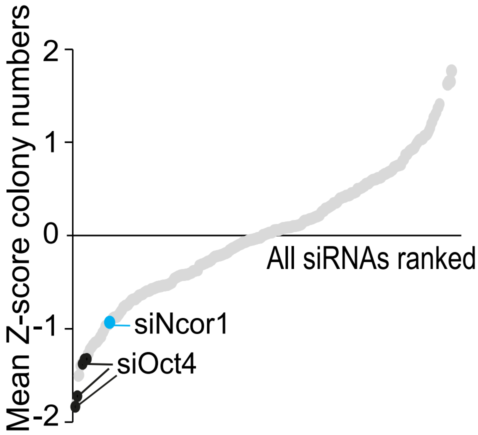


Supplementary Figure 1. Related to Fig.1.

A) High-content images from 9 stitched-fields spanning the whole 96-well, for Ncor1, Oct4 or nt control siRNAs. In green CDH1 staining, SALL4 in magenta and the overlay with DAPI nuclear counterstain. The scale bar represents 500 uM.

B) Derived from the HC-screen (Peñalosa-Ruiz, et.al. 2019), reprogramming efficiencies (colony numbers) from +300 siRNAs, ranked by their Z-score average of four replicates. The black labels correspond to siOct4 and siNcor1(blue) also scores low in colony numbers.

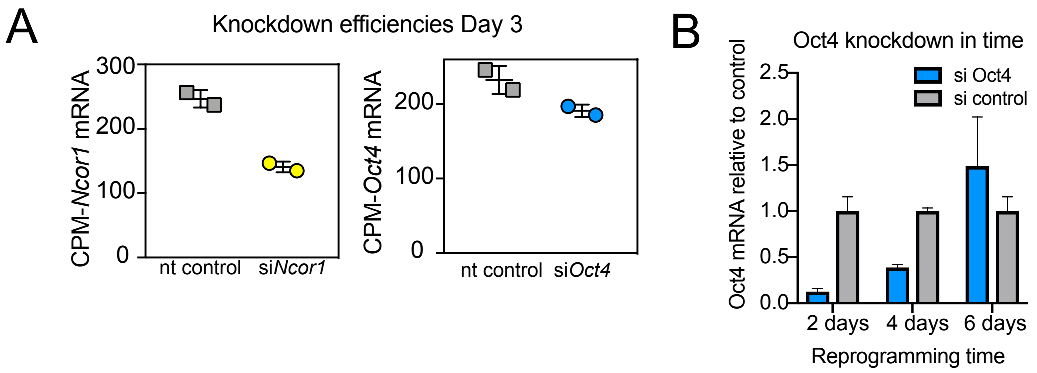


Supplementary Fig. 2. Related to Figure 3.
A) Transcript counts per million reads (CPM) for Ncor1 (yellow) and Oct4 (blue) siRNAs, compared to nt control after RNA-seq, show that the corresponding mRNAs were targeted.
B) Time-course for Oct4 mRNA silencing at different reprogramming time-points. Data was derived from RT-qPCR, where Oct4 mRNA levels were determined relative to tubulin according to the $\Delta\Delta$Ct method. Then, expression values in nt control were set to 1 and Oct4 remaining expression levels in siOct4(blue) were calculated relative to nt values, per time-point. The bars represent the average of two independent transfection replicates +/- Standard Deviations.

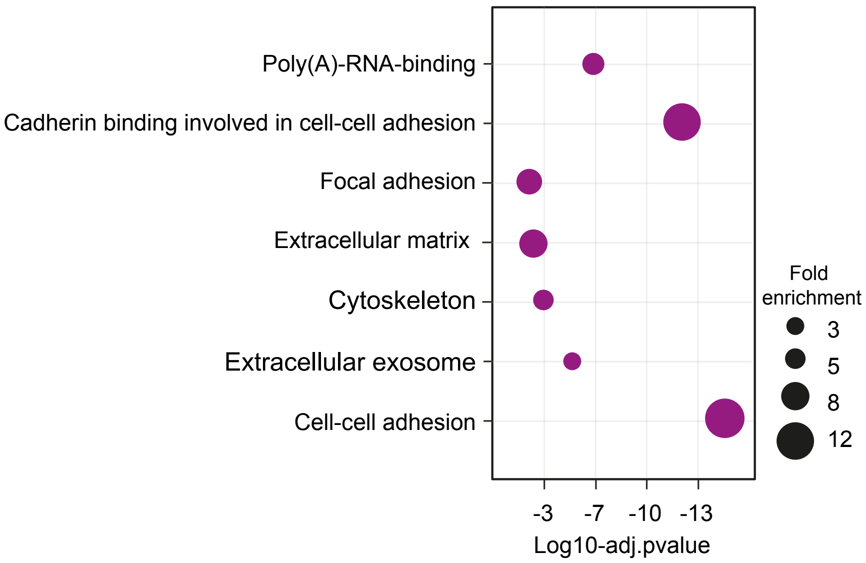


Supplementary Fig.3. Related to Fig.4. Bubble chart depicting GO classification of genes upon Ncor1 knockdown and bound by NCOR1 from ChIP-seq data. The size of the bubble represents the fold enrichment and the x-axis the log10 of the adjusted p-value.This analysis was performed with genes corresponding to Fig.4D.
